# Effect of different derivatization protocols on the calculation of trophic position using amino acids compound-specific stable isotopes

**DOI:** 10.1101/2020.05.12.091785

**Authors:** Stephane Martinez, Maya Lalzer, Eli Shemesh, Shai Einbinder, Beverly Goodman Tchernov, Dan Tchernov

## Abstract

Amino acids compound-specific nitrogen stable isotope (AA-CSIA) is an emerging tool in ecology for understanding trophic system dynamics. While it has been successfully used for several independent studies across a range of environments and study locations, researchers have encountered calculation issues for determining trophic position values. Most studies introduce modifications to the constants of trophic position equation calculations, but then fail to account for the equation variations when comparing the results of separate research studies. The acceptance of this approach is related to the underlying presumption that no addition of the exogenous nitrogen atom occurs in the different methods and, therefore, such variations should not affect the outcome. In this paper, we evaluate the use of the EZfaast amino acid derivatization kit (chloroformate) and compare it to the isotopic results of two other derivatization methods. We highlight new considerations for working with AA-CSIA that might account for some of the variations in the results and lead researchers to modify constants in the equation. This likely requires developing the unique constants per derivatization method in order to be able to compare the trophic position results across different studies.

## 1 Introduction

Traditional methods for calculating the trophic position include stomach content analysis, bulk δ^15^N stable isotope analysis and, more recently, amino acid compound-specific nitrogen isotopic analysis (AA-CSIA). Stomach content analysis provides information on an individual’s prey taxonomy and their relative importance within the food web. However, because it represents only a snapshot in time, this method exhibits bias from a myriad of factors including the proportion of identifiable dietary items, significant numbers of ‘empty stomach’ samples in top predator collections, and varying digestibility of different prey species (i.e. residence time in the stomach). Therefore, when using the stomach content approach, large numbers of samples are needed to correctly evaluate the trophic position (Rindorf and Lewy, 2004), which is both labor-intensive and sometimes impossible. Bulk δ^15^N of whole organisms and their tissues has been used in several ecological studies as an alternative, or in tandem, to assess the trophic position and nitrogen flow in the food web (Yoshii et al., 1999; Post, 2002; Logan and Dodge, 2013; Vander Zanden et al., 2013). This approach maintains the observed relationship between rising δ^15^N (2–4‰) values and higher trophic positions (Minagawa and Wada, 1984). However, the increase in δ^15^N is not constant under all conditions and, therefore, results are only locally applicable and, in most cases are relative to other samples. The value is influenced by the food sources, stressors, consumer physiology, and natural δ^15^N concentration of the surrounding environment. Constraining the nitrogen isotopic baseline, or isotopic composition of primary producers at the base of an ecosystem can be complicated and difficult, or impossible in certain environments (Popp et al., 2007). Thus, researchers were led to seek new methods to determine the trophic position using δ^15^N to better understand baseline values for purposes of interstudy normalization. McClelland and Montoya (2002) were the first to examine AA-CSIA for establishing a trophic position using nitrogen isotope. The research investigated the relationship between lab-cultivated phytoplankton and its consumer, zooplankton. They discovered that the “non-essential” AA glutamic-acid (also known as “trophic” AA) has become “heavier” (richer in δ^15^N) compared to the bulk tissue. The “essential” AA phenylalanine (also known as “source” AA) is inert in terms of trophic position and is not affected by the organism’s position on the food chain. However, it records the δ^15^N signature of the primary producer in the particular food web in question. Since both the isotopic baseline and fractionated information is retained in the nitrogen isotopic composition of AAs, results of AA-CSIA from a single consumer provide both an integrated measurement of trophic position and the δ^15^N value at the base of the food web. This eliminates the need to collect and analyze them independently of the predator species. In order to confirm the applicability of this approach to ecosystem-level studies (rather than species-specific), later studies tested several different macroalgae, phytoplankton, zooplankton gastropods, and fish in the natural environment and lab (Chikaraishi et al., 2007, 2009). It was concluded that due to the different traits of the AAs (glutamic-acid and phenylalanine), the trophic position can be readily calculated TP_Glu/Phe_=((δ^15^N_Glu_–δ^15^N_Phe_–β)/TDF_AA_)+1 without the need to directly measure the primary producers δ^15^N (Chikaraishi et al., 2009; Steffan et al., 2013).

The constant, β, is the difference between the δ^15^N values of glutamic-acid and phenylalanine AAs in primary producers (trophic position 1). The trophic discrimination factor (TDF_AA_) is the average δ^15^N enrichment relative to source AAs per trophic position. When calculating the trophic position based on AA-CSIA, the “β” and “TDF_AA_” are constant to and dependent on the nitrogen source. Chikaraishi et al. (2009) have found that for the marine environment β=3.4±0.9‰, while for terrestrial environment values for β in C3 and C4 plants were -8.4±1.6‰ and -0.4±1.7‰, respectively (Chikaraishi et al., 2010). As for TDF_AA,_ it was thought to be 7.6±1.7‰ for all environments (Chikaraishi et al., 2009, 2010). However, further studies have found that this is not always accurate. Bradley et al. (2015) recalculated the “TDF_AA_” based on a variety of teleost from various trophic positions and found it to be inaccurate in the higher trophic positions, and instead established a value of 5.7±0.3‰. Nielsen et al. (2015) performed similar work and produced calculations of 6.6±1.7‰. McMahon and McCarthy (2016) reviewed the literature and found that the variability is higher (0‰-10‰) and dependent on a range of variables such as nitrogen excretion, diet, and trophic position. Since, presumably, no nitrogen atoms are added in the process of AA derivatization it is thought that using different methods will not influence the final result. Therefore, no comparison has been made between the different methods to check whether this might the reason for the variability.

In this study, we test the influence of different derivatization methods and various locations on the AA isotopic ratio and trophic position *in situ*, in order to determine the influence of these factors on the calculation of trophic position and a better understanding of the used protocols.

## 2 Material and Methods

### 2.1 Sample collection

For comparison between sites, we used samples of fish and algae from the Eastern Mediterranean, Western Mediterranean, and the Red Sea, as well as fish samples from the Indian Ocean (fish only).

### 2.2 Sample preparation

All collected samples were immediately frozen and then lyophilized at the lab prior to the hydrolyzation. Approximately 1.5 mg of fish muscle (between the dorsal fin and the head) and 3-5 mg of algae was acid hydrolyzed in 1 ml of 6 nmol HCl at 150 °C for 75 min (Cowie and Hedges, 1992) under nitrogen atmosphere inside a 4 ml glass vial with PTFE cap. Samples were cooled to room temperature and algae samples were filtered through a 0.22 µ PTFE filter to remove all undissolved particles. The HCl was evaporated under a gentle stream of nitrogen and neutralized twice with 1 ml of ultra-pure water (also evaporated). For chloroformate derivatization, we used an EZfaast amino acid analysis kit, slightly modified by replacing reagent 6 with dichloromethane (DCM) as a solvent. For comparison, herbivorous fish samples were also derivatized following the Metges et al. (1996) protocol for N-Acetyl-n-propyl (NAP)-amino acid derivatization. The third approach from Silfer et al. (1991) used Trifluoroacetic anhydride (TFAA) for the acylation.

For all methods, we injected 1.5 µl in a splitless mode at 250 °C. Helium was used as a carrier gas at a constant flow of 1.5 ml/min for the chloroformate and N-Acetyl-n-propyl; for the TFAA we used 1.1 ml/min. The chloroformate amino acids were separated on a Zebron ZB-50 column (30 m, 0.25 mm, and 0.25 µm) in a Thermo Scientific Trace 1300 Gas chromatographer (GC). Conditions were set to optimize peak separation for the desired amino acids as follows: Initial temperature 110 °C ramped to 240 °C at 8 °C per min and then ramped to 320 °C at 20 °C per min and held for 2.5 min. The N-Acetyl-n-propyl (NAP)-amino acid was separated on the Thermo Scientific TraceGOLD TG-5MS column (30 m, 0.25 mm, and 0.25 µm) in a Thermo Scientific Trace 1300 Gas chromatographer (GC). Conditions were set to optimize peak separation for the desired amino acids as follows: Initial temperature 75 °C ramped to 130 °C at 4 °C per min, held for 2 min, ramped to 180 °C at 5 °C per min, held for 2 min, ramped to 320 °C at 20 °C per min, and held for 1 min.

The TFAA depravities amino acid were separated on the Thermo Scientific TraceGOLD TG-1MS column (30 m, 0.25 mm, and 0.25 µm) in a Thermo Scientific Trace 1300 Gas chromatographer (GC). Conditions were set to optimize peak separation for the desired amino acids as follows: Initial temperature 75 °C held for 1 min ramped to 90 °C at 7.5 °C per min, held for 1.5 min, ramped to 160 °C at 7 °C per min, held for 3.5 min, ramped to 320 °C at 25 °C per min, and held for 2 min. The separated amino acids were split on a MicroChannel Device into two directions, one toward the Thermo Scientific ISQ quadruple for amino acid identification and the second toward the Thermo Scientific Delta V advantage for N_2_ isotope analysis. The ISQ conditions were set to transfer line 310 °C, ion source 240 °C, and scanned in the range 43 to 450 m/z mass range. To define the isotopic ratio of nitrogen, the separated amino acids were combusted in a Thermo scientific GC isolink II at 1000 °C. Before entering to Delta V for the N_2_ analysis, the sample went through a liquid nitrogen cold trap to freeze all other gases. A triplicate was injected from each sample.

### 2.3 Data analysis and corrections

Separated amino acids were purchased from Sigma Aldrich and analyzed with the Geological Survey of Israel’s elemental analyzer isotope ratio mass spectrometer. To extend the nitrogen isotopic range, two certified amino acids (Alanine +43.25‰ and Valine +30.19‰) were purchased from Arndt Schimmelmann (Biogeochemical Laboratories, Indiana University). We used a standard that contains seven amino acids with a known isotopic ratio (Alanine, Valine, Leucine, Isoleucine, Methionine, Glutamic acid, and Phenylalanine) with an isotopic range for the nitrogen of -6.69‰ to +43.25‰. Since nitrogen is not added in the process of derivatization, corrections for nitrogen addition were not required. The standard of amino acids was injected three times after the combustion reactor oxidation and to allow for drift correction, the standard was injected again three times after a maximum of 18 sample injections. Since AAs differ in the presence of heteroatoms and functional groups, this may lead to different combustion efficiencies and, therefore, variation in drift. To compensate for this drift an average of the standard injection from the beginning and the end of the sequence were used. For each sequence, a correction factor was applied based on the linear regression equation of the ratio between the known AA isotopic ratio and the acquired result for the sequence. Stable isotope ratios were expressed in standard δ notation where the standard for nitrogen is atmospheric N_2_ (air).

### 2.4 Trophic Position calculation

The trophic position was calculated from the equation

TP_Glu/Phe_=((δ^15^N_Glu_–δ^15^N_Phe_–β)/TDF_AA_)+1 (Chikaraishi et al., 2009). To examine the influence of the site factor on the trophic position, we re-calculated the values needed for the equation. All samples analyzed were from the eastern Mediterranean Sea. To calculate β, we used 10 different algae (Supplementary Table 1). For TDF_AA_ calculation, we used 17 herbivorous fish (*Siganus rivulatus* and *S. luridus*) (Supplementary Table 2). Calculations yielded the following values: β=-0.36 ±1.49‰ and TDF_AA_=4.54±1.36‰.

## 3 Results

### 3.1 Trophic position of samples from different locations

Samples from different locations (Red Sea, Indian Ocean, and western Mediterranean Sea) were compared for the calculated trophic position against samples from the eastern Mediterranean Sea (Figure 1 and Supplementary Table 3). Trophic position calculations were based on AA-CSIA of nitrogen using the equation we built on the eastern Mediterranean Sea samples. A Kruskal-Wallis rank-sum test was used to determine the significant difference between samples. We did not find any significant differences between samples.

**Figure 1.**
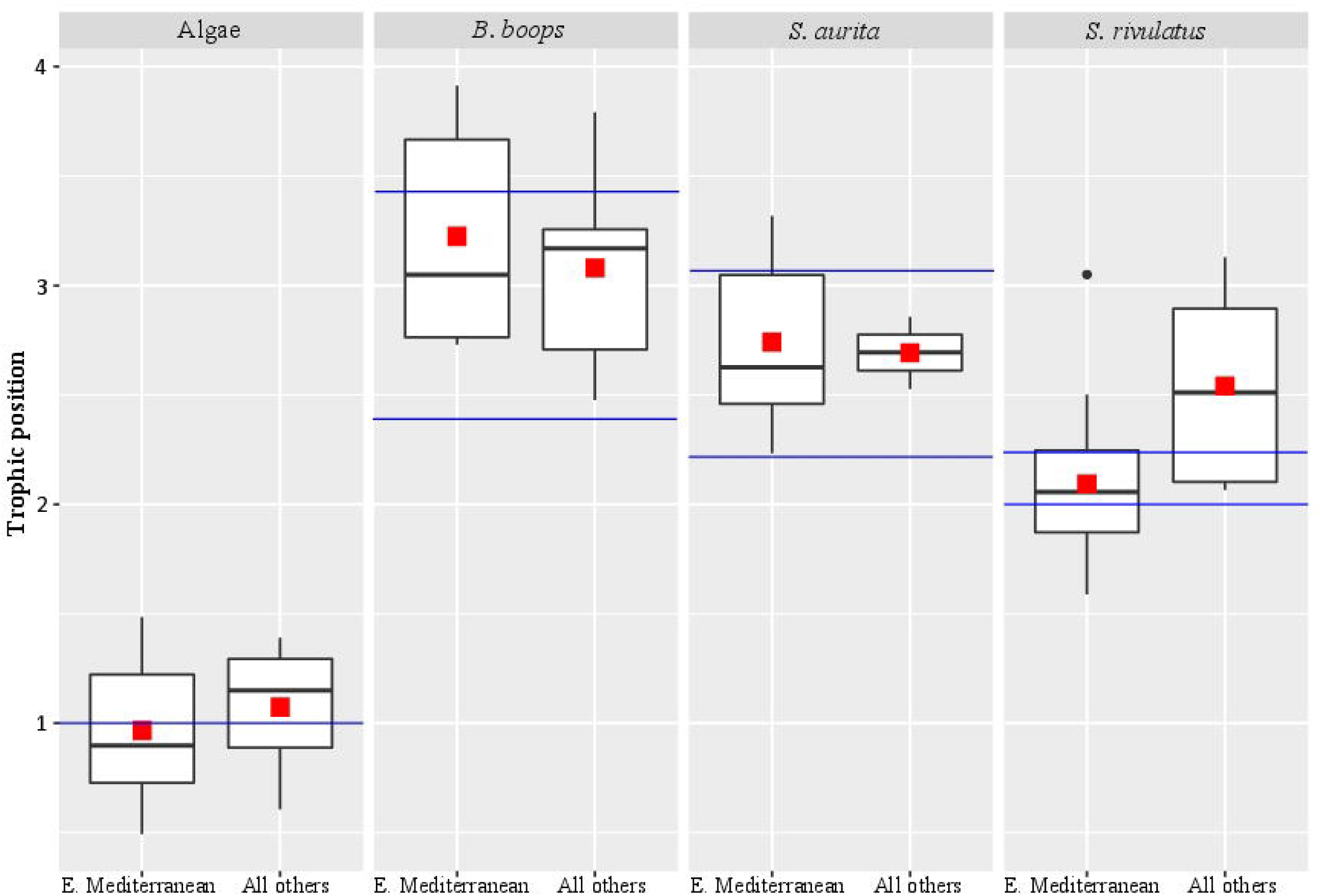
Trophic position of samples from different locations. The red square marks the average trophic position, the black bar in the box marks the median, the black dots are outlier values and the blue bars mark the literature-based trophic position range.

### 3.2 Comparison of the nitrogen isotopic values between different derivatization methods

*S. rivulatus* (n=10) was analyzed using three different methods: chloroformate, NAP (N-Acetyl-n-propyl), and TFAA. We compared Glutamic acid and Phenylalanine δ^15^N (Figure 2 and Supplementary Table 4), the two most commonly used amino acids for calculation of trophic position (Chikaraishi et al., 2009). We applied a Kruskal-Wallis rank-sum test and then adjusted the p-values with the Benjamini-Hochberg method. There was a significant difference in Glutamic acid between Chloroformate to NAP and between Chloroformate to TFAA. In the Phenylalanine, there was a significant difference between Chloroformate to TFAA and between TFAA to NAP. There was no consistency in the shift of the isotopic value between the methods and, therefore, no correction factor can be applied.

**Figure 2.**
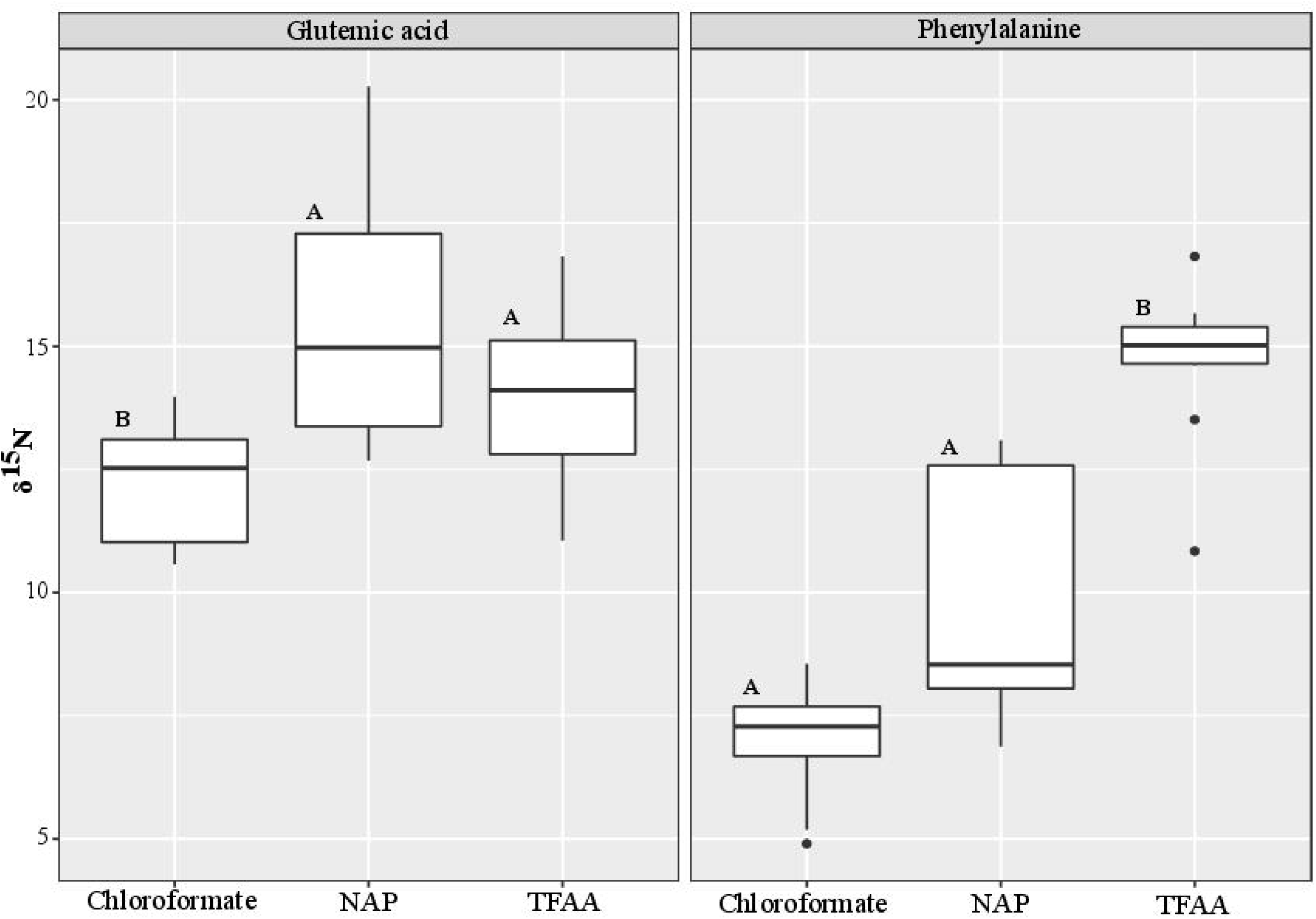
Comparison of the nitrogen isotopic values between different derivatization methods. The black bar in the box marks the median δ^15^N value and the black dots are outlier values. Statistical significance is indicated in bold letters, and p-values are considered for p < 0.05.

## 4 Discussion

In reviewing the literature related to the calculation of the trophic position of teleost using AA-CSIA, it was noted that different studies used different derivatization methods (e.g. Chikaraishi et al. 2009; Bradley et al. 2015; Nuche-Pascual et al. 2018). In all these methods, no correction was made for nitrogen, given that no nitrogen atoms were added in the process. In this study, we used the EZfaast kit, as it is considered to be the easiest, fastest, and safest method to work with; though it is not reported to be used in previous AA-CSIA studies. The isotopic ratio results from the present analysis, however, did not match any known equations in the literature. This research was concentrated on the Eastern Mediterranean Sea, which is ultraoligotrophic and phosphate-limited, even when compared to classic “blue deserts” such as tropical coral reef environments and mid-ocean gyres in the Pacific Ocean (Krom et al., 2010). Because of these conditions, we must distinguish between the potential effect of the method we are using and the unique influence of local environmental effects. To resolve this, we performed our measurements both on the eastern Mediterranean Sea as well as the Red Sea, Indian Ocean, and western Mediterranean Sea samples (Supplementary Table 3). None of the results matched previously reported values using traditional equations. Therefore, we decided, initially, to form our equation based on samples that are readily available locally. For calculating β we used ten different algae which produced a β value of -0.36 ±1.49‰. To calculate the TDF_AA_, we chose two herbivorous species (*S. rivulatus* and *S. luridus*) that, based on literature, has a purely herbivorous diet (Woodland, 1990). From those 17 specimens, we calculated the TDF_AA_ value of 4.54 ± 1.36‰. Using these newly calculated constants, we compared the trophic position of samples from the eastern Mediterranean Sea to samples from the other sources (Red Sea, Indian Ocean, and the Western Mediterranean). We did not find any significant difference between the eastern Mediterranean Sea to the other samples (Figure 1), hence verifying it was not an environmental factor that caused the differential results in trophic position. Also, the trophic position we calculated for *Boops boops* and *Sardinella aurita* samples from all locations are in the range that is reported in the literature (Tsikliras et al., 2005; Bode et al., 2006; Madkour, 2012; Mancinelli et al., 2013; Cresson et al., 2014; Albo-Puigserver et al., 2016). Although not significantly different from the Mediterranean Sea, the trophic position measured for the Red Sea samples of *S. rivulatus*, described as a pure herbivore (TP=2.1±0.1) were slightly higher (TP=2.5±0.5) than reported in the literature (Woodland, 1990). That might be due to the reason that these same species were recently documented eating ctenophores and scyphozoans on top of algae and other invertebrates that are part of the alga biome (Bos et al., 2017; Guy-Haim et al., 2017). Therefore, we conclude that although in many aspects the eastern Mediterranean Sea is a unique environment, the measured variations as compared to other places are not significant enough to impact the trophic position and, therefore, the equation is robust enough to be more broadly applied.

To further validate the equation (in combination with the technique), we tested ten different samples of *S. rivulatus* using three different methods (Chloroformate (EZfaast), NAP (N-Acetyl-n-propyl) and TFAA; Supplementary Table 4). We compared Glutamic acid and Phenylalanine δ^15^N, the two most widely used amino acids for trophic position calculations (Figure 2). Although nitrogen is not added in any of the derivatization protocols, we still observe differences in the isotopic ratios of nitrogen. Our study is in agreement with a previous study by Hofmann et al. (2003) that found differences between the isotopic values of different analytical methods and, therefore, an additional source must be present that causes these isotopic differences. There is a multitude of possible explanations. We can attribute the difference to the impurity of the AA, specifically from a non-AA matrix in the derivatization process or some fatty acids that can also go through the derivatization process alongside the amino acids. Another possibility could be related to different AA extraction efficiencies, variations that will occur due to the differential reaction of derivatized compounds with the combustion reactor on different conditions. In addition, glutamic acid can partially be cyclized into Pyroglutamic acid, or a number of other factors (Castro et al., 1997; Goto et al., 2011; Walsh et al., 2014). The consistency between isotopic ratios within any given protocol, but not between different procedures, emphasizes the importance of applying the correct constant of β and TDF_AA_ per specific protocol in order to conduct interstudy comparisons. Here, we adapted the Ezfasst kit (chloroformate) for quick, safe, and easy analyses of AA-CSIA and provided a robust equation for this protocol that allows for accuracy and precision regardless of the geographic origins of the samples.

## 6 Author Contributions

SM, DT, SE, and ML participated in the design of the study. SM and ES conducted the isotopic analysis. SM, DT, and BG, contributed to the writing and improving of the manuscript. SM and ML participated in the data analysis. All authors contributed and approved the manuscript.

## 8 Acknowledgments

The authors would like to thank Dr Tali Mass, Dr Nir Stern and Dr Christine Ferrier Pages for their assistance in collecting samples at the various sites.

