## Supplementary material for "Effect of different derivatization protocols on the calculation of trophic position using amino acids compound-specific stable isotopes": Table 4

Table 4- Comparison of the herbivore fish *S.rivulatus* nitrogen CSIA of glutamic acid (Glu) and phenylalanine (Phe) using three different derivatization protocols N-Acetyl-n-propyl (NAP), Trifluoroacetic anhydride (TFAA) and chloroformate. For each sample, 3 technical replication were made their average (avg) is displayed and their standard deviation (stdev). The average was corrected (corrected) by a linear equation that was obtained from the standard injection value that was performed before and after each sequence with correspondence to their value before derivatization. The trophic position (TP) was calculated using the constants that we obtained for this work. For more information please see material and methods

| **Method** | **sample ID** | **specie** | **Glu 1** | **Glu 2** | **Glu 3** | **avg** | **stdev** | **corrected** | **Phe 1** | **Phe 2** | **Phe 3** | **avg** | **stdev** | **corrected** | **TP** |
| --- | --- | --- | --- | --- | --- | --- | --- | --- | --- | --- | --- | --- | --- | --- | --- |
| TFAA | Baby1 | S.rivulatus | 15.95 | 15.91 | 15.35 | 15.74 | 0.33 | 12.54 | 14.86 | 13.15 | 15.00 | 14.34 | 1.03 | 10.84 | 1.45 |
| TFAA | Baby2 | S.rivulatus | 16.45 | 16.71 | 16.62 | 16.59 | 0.13 | 13.59 | 17.16 | 17.96 | 17.64 | 17.59 | 0.40 | 14.80 | 0.81 |
| TFAA | Baby3 | S.rivulatus | 14.50 | 14.81 | 14.23 | 14.52 | 0.29 | 11.06 | 16.34 | 17.04 | 16.20 | 16.53 | 0.45 | 13.51 | 0.54 |
| TFAA | Baby4 | S.rivulatus | 15.47 | 15.90 | 15.79 | 15.72 | 0.23 | 12.52 | 17.94 | 18.29 | 18.65 | 18.29 | 0.35 | 15.66 | 0.39 |
| TFAA | Baby5 | S.rivulatus | 16.80 | 16.39 |  | 16.60 | 0.29 | 13.60 | 18.75 | 17.36 |  | 18.06 | 0.98 | 15.37 | 0.69 |
| TFAA | SY62 | S.rivulatus | 18.74 | 18.40 | 18.04 | 18.39 | 0.35 | 14.61 |  | 13.65 | 13.85 | 13.75 | 0.14 | 9.57 | 2.19 |
| TFAA | SY64 | S.rivulatus | 20.15 | 20.67 | 20.48 | 20.43 | 0.27 | 16.82 | 15.92 | 17.63 | 17.87 | 17.14 | 1.06 | 13.26 | 1.86 |
| TFAA | SY65 | S.rivulatus | 19.37 | 18.60 | 19.39 | 19.12 | 0.45 | 15.40 | 17.06 | 17.11 | 17.60 | 17.26 | 0.30 | 13.38 | 1.52 |
| TFAA | SY66 | S.rivulatus | 18.65 | 19.38 | 18.89 | 18.97 | 0.37 | 15.24 | 18.09 | 19.45 | 18.13 | 18.56 | 0.77 | 14.79 | 1.18 |
| TFAA | SY67 | S.rivulatus | 18.92 | 18.53 | 18.12 | 18.52 | 0.40 | 14.75 | 19.72 | 18.92 | 18.37 | 19.00 | 0.68 | 15.27 | 0.96 |
| NAP | Baby1 | S.rivulatus | 17.85 | 18.23 | 18.33 | 18.14 | 0.26 | 13.37 | 13.42 | 13.92 | 13.78 | 13.70 | 0.26 | 8.19 | 2.22 |
| NAP | Baby2 | S.rivulatus |  | 17.36 | 17.72 | 17.54 | 0.26 | 12.68 |  | 12.65 | 12.51 | 12.58 | 0.10 | 6.88 | 2.36 |
| NAP | Baby3 | S.rivulatus |  | 17.49 | 18.55 | 18.02 | 0.75 | 13.24 |  | 13.73 | 13.45 | 13.59 | 0.19 | 8.06 | 2.22 |
| NAP | Baby4 | S.rivulatus | 19.78 | 19.71 | 19.02 | 19.50 | 0.42 | 14.97 | 13.22 | 13.29 | 13.06 | 13.19 | 0.12 | 7.59 | 2.70 |
| NAP | Baby5 | S.rivulatus | 18.79 | 18.98 | 18.78 | 18.85 | 0.11 | 14.21 | 14.06 | 13.98 | 13.97 | 14.00 | 0.05 | 8.54 | 2.33 |
| NAP | SY62 | S.rivulatus | 18.92 | 18.95 | 19.02 | 18.96 | 0.05 | 15.67 | 13.50 | 13.43 | 13.29 | 13.41 | 0.11 | 9.13 | 2.52 |
| NAP | SY64 | S.rivulatus | 22.23 | 23.03 | 23.32 | 22.86 | 0.56 | 20.26 | 16.07 | 16.42 | 17.03 | 16.51 | 0.49 | 12.78 | 2.73 |
| NAP | SY65 | S.rivulatus | 21.62 | 21.75 | 21.52 | 21.63 | 0.11 | 18.81 | 16.04 | 16.69 | 16.28 | 16.34 | 0.33 | 12.58 | 2.45 |
| NAP | SY66 | S.rivulatus | 21.10 | 21.74 | 22.71 | 21.85 | 0.81 | 18.27 | 13.07 | 14.09 | 14.31 | 13.82 | 0.66 | 9.42 | 3.03 |
| NAP | SY67 | S.rivulatus | 20.77 | 20.18 | 20.03 | 20.33 | 0.39 | 17.28 | 16.96 | 16.96 | 16.39 | 16.77 | 0.33 | 13.09 | 2.00 |
| Chloroformate | Sy62 | S.rivulatus | 10.85 | 8.35 | 8.58 | 9.26 | 1.38 | 13.67 | 6.35 | 4.10 | 5.83 | 5.42 | 1.18 | 8.55 | 2.21 |
| Chloroformate | SY64 | S.rivulatus |  | 11.56 | 12.35 | 11.96 | 0.56 | 11.90 |  | 7.08 | 7.42 | 7.25 | 0.24 | 6.93 | 2.17 |
| Chloroformate | SY65 | S.rivulatus | 16.58 | 15.52 | 15.46 | 15.85 | 0.62 | 14.05 | 10.68 | 9.56 | 9.50 | 9.91 | 0.67 | 7.55 | 2.51 |
| Chloroformate | sy66 | S.rivulatus | 14.36 | 12.59 |  | 13.47 | 1.25 | 13.97 | 7.44 | 7.25 | 6.19 | 6.96 | 0.67 | 7.51 | 2.50 |
| Chloroformate | SY67 | S.rivulatus | 12.51 | 13.10 | 13.31 | 12.97 | 0.41 | 12.97 | 6.41 | 8.23 | 8.96 | 7.87 | 1.31 | 7.58 | 2.27 |
| Chloroformate | baby1 | S.rivulatus | 12.31 | 12.13 | 12.07 | 12.17 | 0.13 | 12.00 | 8.32 | 7.97 | 7.97 | 8.09 | 0.20 | 7.84 | 2.00 |
| Chloroformate | Baby2 | S.rivulatus | 12.45 | 12.71 | 12.91 | 12.69 | 0.23 | 9.69 | 8.51 | 8.81 | 9.53 | 8.95 | 0.52 | 5.51 | 2.00 |
| Chloroformate | baby3 | S.rivulatus | 13.23 | 13.16 | 13.70 | 13.36 | 0.29 | 13.22 | 10.51 | 10.37 | 11.25 | 10.71 | 0.47 | 10.51 | 1.68 |
| Chloroformate | Baby4 | S.rivulatus | 14.06 | 14.00 | 13.77 | 13.94 | 0.16 | 11.09 | 11.39 | 10.45 | 10.61 | 10.82 | 0.50 | 7.60 | 1.85 |
| Chloroformate | Baby5 | S.rivulatus | 16.32 | 16.09 | 15.84 | 16.08 | 0.24 | 12.09 | 12.56 | 12.53 | 12.28 | 12.46 | 0.16 | 8.15 | 1.95 |
