## Supplementary material for "Effect of different derivatization protocols on the calculation of trophic position using amino acids compound-specific stable isotopes": Table 3

Table 3- Trophic position calculation from Chloroformate (EZfaast) nitrogen CSIA of glutamic acid (Glu) and phenylalanine (Phe) of different species from the Red Sea, western Mediterranean and Indian ocean compared to the eastern Mediterranean sea For each sample, 3 technical replication were made their average (avg) is displayed and their standard deviation (stdev). The average was corrected (corrected) by a linear equation that was obtained from the standard injection value that was performed before and after each sequence with correspondence to their value before derivatization. The trophic position (TP) was calculated using the constants that we obtained for this work. For more information please see material and methods

| **Location** | **Specie** | **Sample** | **Glu1** | **Glu2** | **Glu3** | **avg** | **stdev** | **corrected** | **Phe1** | **Phe2** | **Phe3** | **avg** | **stdev** | **corrected** | **TP** |
| --- | --- | --- | --- | --- | --- | --- | --- | --- | --- | --- | --- | --- | --- | --- | --- |
| Eastern Mediterranean | alga | alga10 | 3.40 | 4.54 | 3.81 | 3.92 | 0.57 | 3.86 | 2.61 | 2.35 | 1.59 | 2.18 | 0.53 | 2.02 | 1.48 |
| Eastern Mediterranean | alga | alga11 | 3.68 | 3.36 | 2.79 | 3.27 | 0.45 | 3.18 | 5.12 | 4.97 | 3.47 | 4.52 | 0.91 | 4.51 | 0.79 |
| Eastern Mediterranean | alga | alga13 | 4.85 | 4.78 | 2.91 | 4.18 | 1.10 | 5.73 | 2.52 | 3.94 | 2.67 | 3.05 | 0.78 | 4.58 | 1.33 |
| Eastern Mediterranean | alga | alga14 | 3.31 | 5.20 | 3.16 | 3.89 | 1.14 | 5.43 | 5.24 | 4.00 | 4.87 | 4.71 | 0.64 | 6.26 | 0.90 |
| Eastern Mediterranean | alga | alga16 | 4.42 | 4.25 | 3.08 | 3.92 | 0.73 | 4.78 | 6.67 | 7.05 | 5.99 | 6.57 | 0.53 | 7.44 | 0.49 |
| Eastern Mediterranean | alga | alga3 | 4.66 | 4.79 | 4.33 | 4.60 | 0.24 | 6.40 | 5.76 | 6.25 | 5.79 | 5.93 | 0.27 | 8.02 | 0.72 |
| Eastern Mediterranean | alga | alga4 | 1.33 | 1.21 | 0.84 | 1.13 | 0.26 | 4.44 | 1.90 |  | 1.03 | 1.47 | 0.61 | 4.86 | 0.99 |
| Eastern Mediterranean | alga | alga5 | 2.68 |  | 2.80 | 2.74 | 0.09 | 4.91 | 0.56 |  | 1.94 | 1.25 | 0.98 | 3.34 | 1.43 |
| Eastern Mediterranean | alga | alga6 | 5.33 | 6.87 | 5.81 | 6.00 | 0.79 | 7.21 | 4.72 | 6.70 | 6.16 | 5.86 | 1.02 | 7.05 | 1.11 |
| Eastern Mediterranean | alga | 1 |  | 3.77 | 3.39 | 3.58 | 0.27 | 3.09 |  | 5.54 | 5.20 | 5.37 | 0.25 | 4.98 | 0.66 |
| Eastern Mediterranean | alga | 2 | 4.67 | 7.90 | 6.04 | 6.20 | 1.62 | 5.87 | 6.77 | 8.81 | 7.47 | 7.68 | 1.04 | 7.43 | 0.73 |
| Red Sea | alga | 10BZ | -0.89 | -1.62 | -1.29 | -1.27 | 0.36 | -4.38 | -0.62 | -1.15 | -1.16 | -0.98 | 0.31 | -4.07 | 1.05 |
| Red Sea | alga | 10CZ | -0.07 | 0.37 | 0.53 | 0.28 | 0.31 | -1.92 | -0.78 | -0.51 | -0.32 | -0.54 | 0.23 | -2.78 | 1.31 |
| Red Sea | alga | 5-2Z | 0.48 | 1.73 | 1.53 | 1.25 | 0.67 | -2.05 | -0.47 | 0.37 | 0.60 | 0.16 | 0.56 | -3.28 | 1.39 |
| Red Sea | alga | 5-3Z | 1.65 | 2.84 | 1.79 | 2.09 | 0.65 | -1.08 | 1.23 | 2.04 | 1.35 | 1.54 | 0.44 | -1.71 | 1.25 |
| Western Mediterranean | alga | Spain2 | 3.06 | 3.82 | 3.68 | 3.52 | 0.40 | 0.53 | 3.96 | 4.92 | 4.87 | 4.58 | 0.54 | 1.63 | 0.84 |
| Western Mediterranean | alga | Spain3 | 7.45 | 7.13 | 9.70 | 8.09 | 1.40 | 5.28 | 10.14 | 10.07 | 10.25 | 10.15 | 0.09 | 7.43 | 0.61 |
| Western Mediterranean | B.boops | monaco B boops1 |  | 14.46 | 13.69 | 14.07 | 0.55 | 11.72 |  | 3.21 | 2.82 | 3.02 | 0.27 | -0.60 | 3.79 |
| Western Mediterranean | B.boops | monaco B boops2 | 16.75 | 16.13 | 15.74 | 16.20 | 0.51 | 15.55 | 5.57 | 6.33 | 7.32 | 6.41 | 0.88 | 5.66 | 3.26 |
| Western Mediterranean | B.boops | monaco B boops3 | 13.48 | 12.63 | 12.93 | 13.01 | 0.43 | 10.53 | 5.11 | 6.68 | 7.34 | 6.37 | 1.15 | 3.14 | 2.71 |
| Western Mediterranean | B.boops | monaco B boops4 | 15.68 | 14.62 | 14.59 | 14.96 | 0.62 | 15.38 | 9.32 | 8.32 | 8.20 | 8.62 | 0.61 | 9.03 | 2.48 |
| Western Mediterranean | B.boops | monaco B boops5 | 16.67 | 16.42 | 16.52 | 16.53 | 0.12 | 16.95 | 6.57 | 6.79 | 7.74 | 7.03 | 0.62 | 7.45 | 3.17 |
| Eastern Mediterranean | S.aurita | FSH104 | 19.24 | 20.47 | 19.38 | 19.70 | 0.68 | 18.32 | 8.91 | 10.32 | 9.14 | 9.45 | 0.76 | 8.15 | 3.32 |
| Eastern Mediterranean | S.aurita | SY84 | 16.68 | 16.18 | 16.19 | 16.35 | 0.29 | 12.98 | 11.37 | 11.54 | 11.90 | 11.60 | 0.27 | 7.73 | 2.23 |
| Eastern Mediterranean | S.aurita | SY85 | 15.65 | 16.36 | 16.60 | 16.20 | 0.49 | 12.81 | 10.73 | 11.26 | 11.10 | 11.03 | 0.27 | 7.10 | 2.34 |
| Eastern Mediterranean | S.aurita | SY87 | 19.44 | 19.81 | 20.33 | 19.86 | 0.45 | 16.14 | 13.12 | 13.10 | 13.78 | 13.33 | 0.38 | 9.11 | 2.63 |
| Eastern Mediterranean | S.aurita | SY88 | 18.34 | 17.87 | 15.97 | 17.39 | 1.25 | 13.48 | 11.44 | 10.81 | 10.91 | 11.06 | 0.34 | 6.66 | 2.58 |
| Eastern Mediterranean | S.aurita | Tr1 | 20.46 | 20.98 | 21.18 | 20.87 | 0.37 | 16.70 | 12.83 | 13.05 | 13.16 | 13.01 | 0.17 | 7.53 | 3.10 |
| Eastern Mediterranean | S.aurita | Tr2 | 18.94 | 18.55 | 17.52 | 18.34 | 0.73 | 13.74 | 11.12 | 10.89 | 10.54 | 10.85 | 0.29 | 5.01 | 3.00 |
| Indian ocean | S.aurita | N14 | 15.61 | 17.48 | 15.68 | 16.26 | 1.06 | 13.64 | 8.42 | 9.84 | 8.87 | 9.04 | 0.73 | 6.30 | 2.70 |
| Indian ocean | S.aurita | N15 | 16.64 | 15.26 | 16.58 | 16.16 | 0.78 | 13.54 | 10.23 | 9.05 | 9.78 | 9.69 | 0.60 | 6.95 | 2.53 |
| Indian ocean | S.aurita | N16 | 15.45 | 15.82 | 15.86 | 15.71 | 0.23 | 13.66 | 7.59 | 8.14 | 8.63 | 8.12 | 0.52 | 5.59 | 2.86 |
| Eastern Mediterranean | S.rivulatus | baby1 | 12.31 | 12.13 | 12.07 | 12.17 | 0.13 | 12.00 | 8.32 | 7.97 | 7.97 | 8.09 | 0.20 | 7.84 | 2.00 |
| Eastern Mediterranean | S.rivulatus | Baby2 | 12.45 | 12.71 | 12.91 | 12.69 | 0.23 | 9.69 | 8.51 | 8.81 | 9.53 | 8.95 | 0.52 | 5.51 | 2.00 |
| Eastern Mediterranean | S.rivulatus | baby3 | 13.23 | 13.16 | 13.70 | 13.36 | 0.29 | 13.22 | 10.51 | 10.37 | 11.25 | 10.71 | 0.47 | 10.51 | 1.68 |
| Eastern Mediterranean | S.rivulatus | Baby4 | 14.06 | 14.00 | 13.77 | 13.94 | 0.16 | 11.09 | 11.39 | 10.45 | 10.61 | 10.82 | 0.50 | 7.60 | 1.85 |
| Eastern Mediterranean | S.rivulatus | Baby5 | 16.32 | 16.09 | 15.84 | 16.08 | 0.24 | 12.09 | 12.56 | 12.53 | 12.28 | 12.46 | 0.16 | 8.15 | 1.95 |
| Eastern Mediterranean | S.rivulatus | siganus1 | 11.53 | 10.94 |  | 11.23 | 0.42 | 11.45 | 6.99 | 7.84 | 11.06 | 8.63 | 2.15 | 8.65 | 1.70 |
| Eastern Mediterranean | S.rivulatus | siganus2 | 8.64 | 8.84 | 7.24 | 8.24 | 0.87 | 8.03 | 2.62 | 6.40 | 2.41 | 3.81 | 2.24 | 3.33 | 2.11 |
| Eastern Mediterranean | S.rivulatus | siganus3 | 10.03 | 8.91 | 10.01 | 9.65 | 0.64 | 9.65 | 7.39 | 6.79 | 8.19 | 7.46 | 0.70 | 7.34 | 1.59 |
| Eastern Mediterranean | S.rivulatus | Sy62 | 10.85 | 8.35 | 8.58 | 9.26 | 1.38 | 13.67 | 6.35 | 4.10 | 5.83 | 5.42 | 1.18 | 8.55 | 2.21 |
| Eastern Mediterranean | S.rivulatus | SY63 | 15.37 | 12.02 | 15.38 | 14.26 | 1.94 | 20.82 | 8.68 | 5.66 | 6.90 | 7.08 | 1.52 | 11.87 | 3.05 |
| Eastern Mediterranean | S.rivulatus | SY64 |  | 11.56 | 12.35 | 11.96 | 0.56 | 11.90 |  | 7.08 | 7.42 | 7.25 | 0.24 | 6.93 | 2.17 |
| Eastern Mediterranean | S.rivulatus | SY65 | 16.58 | 15.52 | 15.46 | 15.85 | 0.62 | 14.05 | 10.68 | 9.56 | 9.50 | 9.91 | 0.67 | 7.55 | 2.5 |
| Eastern Mediterranean | S.rivulatus | sy66 | 14.36 | 12.59 |  | 13.47 | 1.25 | 13.97 | 7.44 | 7.25 | 6.19 | 6.96 | 0.67 | 7.51 | 2.50 |
| Eastern Mediterranean | S.rivulatus | SY67 | 12.51 | 13.10 | 13.31 | 12.97 | 0.41 | 12.97 | 6.41 | 8.23 | 8.96 | 7.87 | 1.31 | 7.58 | 2.27 |
| Red Sea | S.rivulatus | EY03 | 12.57 | 13.02 | 13.04 | 12.88 | 0.26 | 9.80 | 4.50 | 5.14 | 5.27 | 4.97 | 0.41 | 1.55 | 2.90 |
| Red Sea | S.rivulatus | EY04 | 10.31 | 8.86 | 9.83 | 9.66 | 0.74 | 6.45 | 5.86 | 4.76 | 5.48 | 5.37 | 0.56 | 1.97 | 2.07 |
| Red Sea | S.rivulatus | EY05 | 12.90 | 13.31 | 13.73 | 13.32 | 0.41 | 10.28 | 5.23 | 3.85 | 4.34 | 4.47 | 0.70 | 0.97 | 3.13 |
| Red Sea | S.rivulatus | EY06 | 10.69 | 8.52 | 9.56 | 9.59 | 1.08 | 6.36 | 5.97 | 4.07 | 5.46 | 5.17 | 0.98 | 1.70 | 2.10 |
