## Supplementary material for "Effect of different derivatization protocols on the calculation of trophic position using amino acids compound-specific stable isotopes": Table 2

Table 2- Calculation of the TDF_AA_ correction factor for the trophic position equation using Chloroformate (EZfaast) nitrogen CSIA of glutamic acid (Glu) and phenylalanine (Phe) of the herbivore fish *S.luridus* and *S.rivulatus*. For each sample, 3 technical replication were made their average (avg) is displayed and their standard deviation (stdev). The average was corrected (corrected) by a linear equation that was obtained from the standard injection value that was performed before and after each sequence with correspondence to their value before derivatization. The trophic position (TP) was calculated using the constants that we obtained for this work. For more information please see material and methods

| **Sample** | **Specie** | **Glu1** | **Glu2** | **Glu3** | **avg** | **stdev** | **corrected** | **Phe1** | **Phe2** | **Phe3** | **avg** | **stdev** | **corrected** | **TP** | **TDF correction** |
| --- | --- | --- | --- | --- | --- | --- | --- | --- | --- | --- | --- | --- | --- | --- | --- |
| siganus3 | *S.rivulatus* | 10.03 | 8.91 | 10.01 | 9.65 | 0.64 | 9.65 | 7.39 | 6.79 | 8.19 | 7.46 | 0.70 | 7.34 | 1.43 | 2.68 |
| baby3 | *S.rivulatus* | 13.23 | 13.16 | 13.70 | 13.36 | 0.29 | 13.22 | 10.51 | 10.37 | 11.25 | 10.71 | 0.47 | 10.51 | 1.68 | 3.07 |
| siganus1 | *S.rivulatus* | 11.53 | 10.94 |  | 11.23 | 0.42 | 11.45 | 6.99 | 7.84 | 11.06 | 8.63 | 2.15 | 8.65 | 1.54 | 3.16 |
| baby5 | *S.rivulatus* | 11.44 | 11.32 | 10.94 | 11.24 | 0.26 | 10.13 | 7.66 | 7.72 | 7.83 | 7.74 | 0.08 | 6.40 | 1.90 | 4.09 |
| baby2 | *S.rivulatus* | 12.24 | 11.61 | 11.98 | 11.94 | 0.32 | 11.77 | 7.82 | 7.89 | 8.36 | 8.02 | 0.30 | 7.77 | 1.96 | 4.36 |
| baby1 | *S.rivulatus* | 12.31 | 12.13 | 12.07 | 12.17 | 0.13 | 12.00 | 8.32 | 7.97 | 7.97 | 8.09 | 0.20 | 7.84 | 2.00 | 4.52 |
| siganus2 | *S.rivulatus* | 8.64 | 8.84 | 7.24 | 8.24 | 0.87 | 8.03 | 2.62 | 6.40 | 2.41 | 3.81 | 2.24 | 3.33 | 1.95 | 5.06 |
| SY64 | *S.rivulatus* |  | 11.56 | 12.35 | 11.96 | 0.56 | 11.90 |  | 7.08 | 7.42 | 7.25 | 0.24 | 6.93 | 2.01 | 5.33 |
| Sy62 | *S.rivulatus* | 10.85 | 8.35 | 8.58 | 9.26 | 1.38 | 13.67 | 6.35 | 4.10 | 5.83 | 5.42 | 1.18 | 8.55 | 2.05 | 5.48 |
| SY67 | *S.rivulatus* | 12.51 | 13.10 | 13.31 | 12.97 | 0.41 | 12.97 | 6.41 | 8.23 | 8.96 | 7.87 | 1.31 | 7.58 | 2.11 | 5.75 |
| baby4 | *S.rivulatus* | 13.97 | 13.32 | 13.79 | 13.69 | 0.33 | 12.76 | 8.96 | 7.61 | 8.21 | 8.26 | 0.68 | 6.96 | 2.36 | 6.16 |
| sy66 | *S.rivulatus* | 14.36 | 12.59 |  | 13.47 | 1.25 | 13.97 | 7.44 | 7.25 | 6.19 | 6.96 | 0.67 | 7.51 | 2.50 | 6.82 |
| SY70 | *S.luridus* | 9.16 | 8.03 | 8.36 | 8.52 | 0.58 | 9.45 | 7.23 | 6.31 | 6.77 | 6.77 | 0.46 | 7.68 | 1.47 | 2.13 |
| SY68 | *S.luridus* | 6.28 | 8.67 | 8.67 | 7.87 | 1.38 | 8.80 | 2.71 | 3.80 | 4.56 | 3.69 | 0.93 | 4.57 | 2.01 | 4.60 |
| SY69 | *S.luridus* | 7.97 | 7.80 | 8.23 | 8.00 | 0.22 | 8.93 | 4.00 | 3.66 | 4.07 | 3.91 | 0.22 | 4.79 | 1.99 | 4.50 |
| SY63 | *S.rivulatus* | 16.01 | 16.42 |  | 16.22 | 0.29 | 14.86 | 11.38 | 12.32 |  | 11.85 | 0.66 | 10.14 | 2.12 | 5.08 |
| SY65 | *S.rivulatus* | 16.58 | 15.52 | 15.46 | 15.85 | 0.62 | 14.05 | 10.68 | 9.56 | 9.50 | 9.91 | 0.67 | 7.55 | 2.51 | 6.87 |
| **avg TDF correction** | | **stdev TDF** | |  |  |  |  |  |  |  |  |  |  |  |  |
| *4.54* | | 1.36 | |  |  |  |  |  |  |  |  |  |  |  |  |
